# Microbial retention and resistances in stormwater quality improvement devices treating road runoff

**DOI:** 10.1101/2021.01.12.426166

**Authors:** Renato Liguori, Steffen H. Rommel, Johan Bengtsson-Palme, Brigitte Helmreich, Christian Wurzbacher

**Author notes:** Corresponding to: C. Wurzbacher, R. Liguori.

## Abstract

Current knowledge about the microbial communities that occur in in urban road runoff is scarce. Road runoff of trafficked roads can be heavily polluted and is treated by stormwater quality improvement devices (SQIDs). However, microbes may influence the treatment process of these devices or could lead to stress resistant opportunistic microbial strains. In this study, the microbial community in the influent, effluent and the filter materials for the removal of dissolved heavy metals of two different SQIDs were analyzed to determine the microbial load, retention, composition, and mobile resistance genes. Although the microbes were replaced by new taxa in the effluent, there was no major retention of microbial genera. Further, the bacterial abundance of the SQIDs effluent was relatively stable over time. The heavy metal content correlated with *intl1* and with microbial genera. The filter media itself was enriched with *Intl1* gene cassettes, carrying several heavy metal and multidrug resistance genes (e.g. *czrA*, *czcA, silP*, *mexW* and *mexI*), indicating that this is a hot spot for horizontal gene transfer. Overall, the results shed light on road runoff microbial communities, and pointed to distinct bacterial communities within the SQIDs, which subsequently influence the microbial community and the genes released with the treated water.

## 1. INTRODUCTION

Industrialization and technological advancement have put an increasing burden on the environment by releasing large quantities of hazardous contaminants inflicting serious damage on the ecosystem. Traffic area runoff is widely recognized as a major transport vector of pollutants released in the urban environment, as it collects precipitation and snowmelt-related discharges of mostly impervious surfaces (e.g. sidewalks, parking lots, feeder streets, major roads, and highways). The majority of pollution caused by traffic area runoff originates from vehicle brake emissions, tire wear, lubricating oil and grease, pet waste and atmospheric deposition on the road surface (1–3). The chemical quality of traffic area runoff has been analyzed and indicated the presence of different contaminants including heavy metals, polycyclic aromatic hydrocarbons, polychlorinated biphenyls (PCB), and other organics (4, 5). Heavy metals in traffic area runoff continue to create serious global health concerns, as they persist in suspended particulate matter and have the ability to travel long distances through water-air-soil systems with subsequent risks to human health (6). The awareness of stormwater runoff pollution and increasing concern about its impacts on the environment, has led to the development of stormwater control measures (SCMs) for pollution control and contaminant retention from urban road runoff. One SCM to minimize the contaminant emissions to the environment is the usage of sustainable urban drainage systems (7, 8). These include decentralized technical systems, referred as stormwater quality improvement devices (SQIDs) or manufactured treatment devices that treat stormwater runoff with a comparably low footprint, and are particularly suitable in dense urban environments (7). SCMs have historically been constructed for pollutants and nutrient reduction from different environments (9, 10). Nevertheless, many studies on the impact of SCMs have also evaluated their function for removal of bacteria, in particular filter-based bioretention systems have shown fecal bacteria removal efficiencies between 50% and 70% with significant difference between inflow and outflow concentrations (11, 12). Pathogenic bacteria, viruses and protozoa can be found in runoff (13, 14) and are transported to surface waters through sewer overflows, representing one of the major human health risks (15, 16).

While the microbial load of sewer overflows has gained considerable attention, microbes of traffic area runoff in general are scarcely investigated with only few exceptions. Due to the heavy pollution and harsh conditions, traffic areas and their runoff can be classified as an extreme environment. Early research looked mainly at microbial communities found in sediments of infiltration basins (17), on the effect of de-icing salts (18, 19), the biofilm of road gutters (20), or on the denitrification potential of road runoff receiving biofilters (21). More comprehensive analysis on the microbiology of road runoff are missing and most of the microbes and their community functioning in this polluted environment remain unknown, despite we can assume that they will be of relevance for the receiving water bodies (groundwater and surface waters) (22, 23). Furthermore, understanding how antibiotic resistance genes (ARGs) are distributed along road runoff treatment processes is important due to their potential public health risk. The location of ARGs on mobile genetic elements, such as integrons (i.e. *intl1, sul1*) makes the horizontal transfer of antibiotic resistance possible and easy to achieve among bacteria with same or diverse origins. Previous studies assessed the presence of Intl1 in the genome of bacterial isolates from wastewater and drinking water treatment plants (24, 25) showing a high possibility of horizontal gene transfer of ARGs within these systems. Therefore, an investigation of these microbial communities can provide insights into adaptations of microbial communities to factors such as pH, contaminants and heavy metals (26, 27), and shed light on potential microbial risks.

This study is a pioneer study on the microbial community composition and its anthropogenic signatures in the form of class I integron gene casettes (*intl1*) of road runoff effluents and the filter materials of two SQIDs along a heavily trafficked urban road in Munich, Germany. We collected water samples for over seven months, and sampled the filter media of the SQIDs. The aims of this study were to: (I) identify the major taxa of road runoff and treated effluent and (II) identify microbial risk factors associated with the mobile genetic element *intl1*. This establishes the basis for evaluating the microbial relevance in road runoff, its impact on receiving water bodies and how this impact may be modulated by current treatment systems.

## 2. MATERIALS AND METHODS

### 2.1 Study site

In this study we monitored two different SQIDs (D1, D2, Figure S1) from a heavily trafficked road in Munich (Germany, 48°10′47″N 11°32′25″E) with an annual average daily traffic of 24,000 vehicles per day. Device D1 and D2 are pre-manufactured SQIDs (SediSubstrator XL 600/12, Fränkische Rohrwerke Gebr. Kirchner GmbH & Co. KG, Germany; Drainfix Clean 300, Hauraton GmbH & Co. KG, Germany). Both SQIDs were installed at the same time and drained road runoff from catchments in close proximity, which consequently showed the same runoff properties. The main difference between the devices were that D1 used a primary sedimentation stage and downstream media filtration stage using an iron-based filter medium with lignite addition and the filter medium was permanently submerged, while D2 used direct filtration with a carbonate containing sand, which ran dry after each rain event. After the treatment, the water was percolated into the groundwater. The catchment areas of the devices were 1660 m^2^ for D1 and 165 m^2^ for D2.

### 2.2 Sampling and characterization

Water samples before (n=21) and after SQID treatment (n=20) were withdrawn during a seven-month time frame starting from April to October 2019, in order to evaluate different seasonal change (spring, n=19; summer, n=13; autumn, n=9). Three different types of water samples were collected based on the position of sampling: Influent (I): inflow of road runoff to the SQIDs; Effluent after sedimentation and adsorption (ESA): effluent of SQID D1; Effluent of Filtration (EF) filtrated water samples of the SQID D2. The samples were withdrawn volume proportionally using automatic samplers (WS 316, WaterSam, Balingen, Germany). Permanent flow measurement using electro-magnetic flow meters (Krohne Optiflux 2300 C or 1300 C, Krohne IFC 300 C, DN250 for D1, DN25 for D2) enabled the volume proportional sampling. Sampling was triggered, if flow exceeded for 1 min 0.4 L/(s·ha) and stopped if flow was below the threshold value for 15 min. Discrete samples (400 mL) were withdrawn after approximately 0.2 L/m^2^ runoff volume each. The discrete samples of each sampling point and runoff event were merged in composite samples for further analysis. The number of discrete samples per sampling point and runoff event depended on the runoff volume and ranged from 2 to 62. The samples were kept in coolers at 4±1°C and transported to the lab within 60 h. Electric conductivity (EC) and pH of the samples were analyzed following the standard methods 2510 B and 4500-H+, respectively (28). Total concentrations of chromium (Cr), copper (Cu), nickel (Ni), lead (Pb) and zinc (Zn) were determined after aqua regia digestion according to EN ISO 15587-1:2002. Cd, Cu, Ni, and Pb were analyzed using ICP-MS (NexION 300D, Perkin Elmer, Waltham, USA). The other elements were analyzed using ICP-OES (DIN EN ISO 11885, Ultima II, Horiba Jobin Yvon, Kyoto, Japan). The limits of quantification (LOQs) were 1.0, 0.1, 0.4, 0.1, and 2.0 µg/L for Cr, Cu, Ni, Pb, and Zn, respectively. Dissolved concentrations of Cr, Cu, Ni, Pb, and Zn were analyzed for a subset of samples after filtration using syringe filter (0.45 µm, PES, VWR International, Darmstadt, Germany). The LOQs of the dissolved Cr, Cu, Ni, Pb, and Zn concentrations were 2.0, 1.0, 1.0, 1.0, and 1.0, µg/L respectively.

Filter material samples (> 500 g) were withdrawn from the SQIDs after approximately 2.75 years of operation, labeled FD1 for SQID D1 and FD2 for SQID D2. The surface layer (0-5 cm) in flow direction of the filter materials, which commonly contain most of the contaminants (29–31), was sampled using ethanol cleaned plastic spatulae and a stainless-steel soil sampler. In addition, we took samples for the microbial community analysis in the middle (5-10 cm) and deepest layer (10-15 cm) in the flow direction of the filter materials. The content of Cr, Cu, Ni, Pb, and Zn in the filter media were analyzed after inverse aqua regia digestion adapted from DIN EN 13346:2001 with a HNO_3_:HCl ratio of 3:1 using the aforementioned ICP-MS and ICP-OES devices. The LOQs of Cr, Cu, Ni, Pb, Zn in the filter media were 5.0, 5.0, 2.0, 10.0, and 1.0 mg/kg, respectively.

All analysis results for water and filter media samples below LOQ were substituted by the respective LOQ value as a conservative estimate.

To assess the overall pollution level of the water samples, a water pollution index (WPI_GFS_) was determined based on the German insignificance threshold values for evaluation of locally restricted groundwater pollution (*Geringfügigkeitsschwellenwerte, Table S1*), which are used to evaluate if a negative anthropogenic effect on groundwater quality is present, following eq. 1. This method is adapted from Bartlett et al., 2012

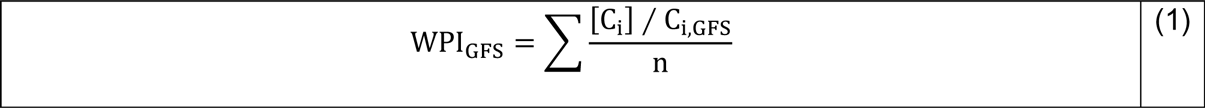

where [C_i_] is the concentration of the substance i present in the sample, C_i,GFS_ is the minor threshold value of substance i, and n is the number of analyzed substances. The heavy metals Cr, Cu, Ni, Pb, and Zn were considered in this analysis.

**Table 1.**
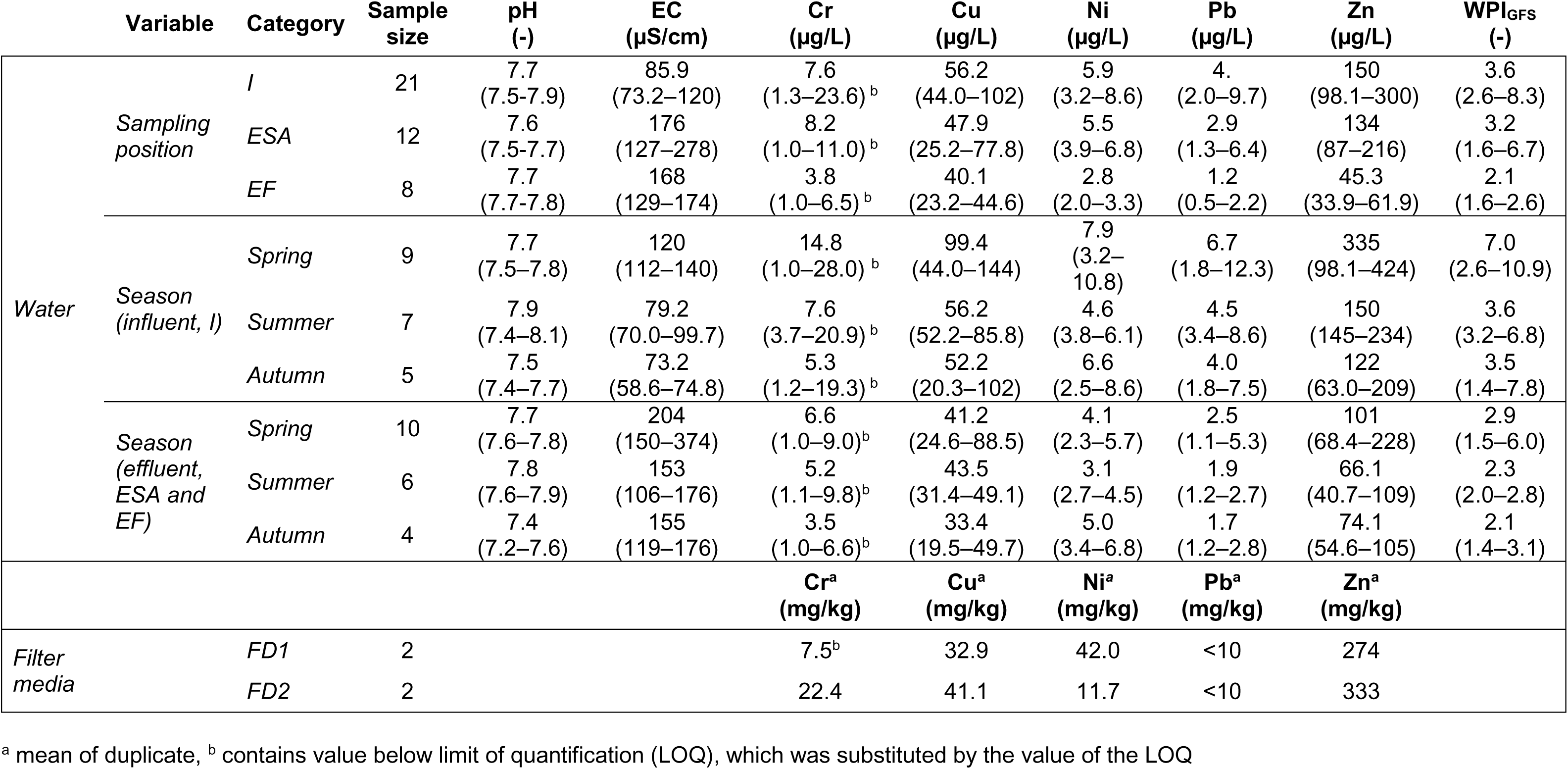
Chemical analysis of the water and filter media samples, reported as median (25%–75%); total concentrations of Cr, Cu, Ni, Pb, and Zn are presented. EC: electric conductivity, and WPI_GFS_: water pollution index. The WPI_GFS_ was added to summarize the contamination level of the samples. I: Influent; ESA: Effluent of device D1; FD1: Filter material of D1; FD2: filter material of D2; EF: Effluent of device D2.

### 2.3 DNA Extraction and 16S rRNA Gene Amplicon Sequencing

Water samples collected from the different devices were centrifuged at 5000 rpm for 10 minutes and the pellets were stored at −20 °C, while the filter media samples were directly stored at −20°C until DNA extraction. The DNA was extracted using the FastDNA Spin Kit for Soil (MP Biomedicals, Solon, USA), following the manufacturers protocol. The DNA concentration of the individual extracts was quantified by using the dsDNA Broad Range Assay kit (DeNovix, Wilmington, USA), following the manufacturers protocol, then stored at −80 °C until sequencing. The 16S rRNA gene amplicon sequencing was performed at ZIEL using the primers 341F/806R targeting mainly bacteria (Institute for Food & Health at Technical University of Munich, Germany). All the data are generated using a MiSeq sequencer (Illumina technology, v3 chemistry) following the protocol of (34)

### 2.4 Data Analysis and Quality Control

All 16S rRNA gene amplicons were processed using the open-source bioinformatic pipeline DADA2 (version1.14.1, Callahan et al., 2016) for R (version 3.6.0) (36). Demultiplexing and quality filtering were carried out in DADA2 using customized settings (truncLen=c(290, 200), trimLeft = c(14, 12), maxN=0, maxEE=c(2, 6)) after the removal of the primers sequence. Error rates were subsequently estimated from a set of subsampled reads (1 million random reads), and chimeric sequences were identified and removed from the demultiplexed reads. The exact amplicon sequence variants (ASVs) were taxonomically classified with a naïve Bayesian classifier using the Silva v. 138 training set (https://benjjneb.github.io/dada2/training.html, accessed August 2020). Negative controls, (i.e. Nuclease-free water) were included at every step of processing, from DNA extraction through the library preparation. A subset of control samples were sequenced in sequencing runs to verify that methodological errors did not impact results. Samples that shared dominant taxa with negative controls were removed from the dataset.

### 2.5 Quantitative Polymerase Chain Reaction (qPCR)

A quantitative Polymerase Chain Reaction (qPCR) protocol was performed to quantify the number of *16S* rRNA and *intl1* gene copies within samples. *16S* rRNA gene was amplified by *16S* Forward 5’-GACTCCTACGGGAGGCWGCAG-3’; *16S* Reverse: 5’-GTATTACCGCGGCTGCTGG-3’ (37). *Intl1* was amplified with the *intl1* primers from (38). The qPCR for 16S rRNA was carried out with a reaction mixture containing 10.5 µL GoTaq® qPCR Master Mix (2X) (Promega, Madison, USA), 0.2 µM of each primer, 7.5 µL nuclease free water and 1 µL of template for a total volume of 21 µL. The 16S rRNA qPCR program consisted of 2 min at 95 °C, 40 cycles with 5 s at 95 °C, 30 s at 60 °C, while for *intl1* the program was 4 min at 95 °C, 40 cycles with 10 sec at 95 °C, 45 s at 64 °C. Both were performed using the CFX96 thermocycler (BioRad, Hercules, USA). Calibration curves for *intl1* were obtained using serial dilutions of a purified PCR products (by NGSBeads, Steinbrenner, Wiesenbach, Germany, following the manual) derived from wastewater. Calibration curves for 16S rRNA were obtained by serial dilutions of a linearized plasmid (pGEM-T easy, Invitrogen, Carlsbad, USA) carrying a single amplicon variant. Specificity of PCR reactions was checked by melt curves, and potential false positives were removed. All samples were analyzed in technical duplicates to obtain final copy numbers per sample by averaging.

### 2.6 *Intl1* gene cassette sequencing

Genomic DNA from three effluent water and two filter material samples of the devices D1 and D2, was used for characterization of class 1 integron gene cassette arrays (*intl1*). The cassette arrays of Tn*402*-associated class 1 integrons were amplified using the primers HS458 and HS459 (39). These primers target the integron recombination site, and the 3’ end of the cassette array, which normally terminates in the *qacEΔ*/*sul1* gene fusion. Sequencing of HS458/459 PCR products can thus recover resistance determinants. The library collection was carried out with the following cycling program: 94 °C for 3 min; 94 °C for 30 s, 55 °C for 30 sec, 72 °C 1 min 30 s for 35 cycles and 72 °C for 5 min. Amplicons were pooled and sequencing was performed using MinION (Oxford Nanopores Technologies, Oxford, UK) using the LSK-109 library preparation according to the manufacturers recommendations and a Flongle flow cell generating 622,526 reads with the high accuracy basecalling mode (MinKnow version 19.10.1).

### 2.7 *Intl1* gene analysis

MinION fastq reads were converted to FASTA format using pefcon (part of the PETKit,, Bengtsson-Palme, 2012) and translated into all six reading frames using the EMBOSS transeq tool (40), options “-trim -clean -frame 6”. Resistance genes were identified using local Usearch (41) against the ResFinder (42), FARME (43) and BacMet experimentally confirmed (44) databases with 70% identity threshold (options “-id 0.7 -blast6out out.blastp -evalue 0.001”). Prior to this search, the FARME database was filtered to contain only actual antibiotic resistance protein sequence, following the protocol in (45). A similar approach was taken to identify markers for mobile genetic elements, using the MGEDB as reference (46) (usearch local options “-id 0.7 -blast6out out.blastp -strand both”). The six-frame translations were also scanned against Pfam (47) using HMMER (using defined trusted thresholds, the “--cut_tc” option). All annotations were added to a FARAO annotation database (48). Lists of annotated integron regions were then produced by querying the FARAO database with different criteria.

### 2.8 Statistical Analysis

Statistical analysis of the microbial community composition was performed by converting the ASV table produced by DADA2 into phyloseq objects using the “phyloseq” package (v.1.24.2) in R (v 3.6.0) (36, 49). The microbial diversity indices were analyzed using the “vegan” and “betapart” package from CRAN (50, 51). The Shannon index was used for the alpha diversity while ASV richness estimate was determined by rarefying the amplicon dataset to the smallest sample (3538). Kruskal-Wallis was used to test significant differences between experimental conditions. Differential abundance analysis of taxa to identify the removal/replacement of microbes before and after the SQIDs was performed by DESeq2 (v 1.29.5) (52). To gain insight about the overall microbial retention exerted by the SQIDs, we partitioned the β-diversity into two components: turnover (β-sim) and nestedness (β-ness) (51). Multivariate statistics were investigated with generalized linear models (GLMs) for multivariate abundance data using the mvabund package (53). Predictive models were fitted using “negative.binomial” family, often being appropriate for count data, with the mean–variance function tending to be quadratic rather than linear. Non-metric multi-dimensional Scaling (NMDS) was used to visualize the microbial community composition and how it aligned with different variables (heavy metals, *intl1*, sample type). The *intl1* data were further normalized by the 16S rRNA copy numbers. The qPCR data (16S rRNA*, intl1*) were log-transformed prior to statistical analysis. We used the BioEnv approach (54) to examine the best subset of environmental variables, correlating with community dissimilarities. In addition, to explore the correlation between microbial community’s relative abundance, heavy metals, and *intl1* gene abundances, Spearman correlations were calculated. To test if the heavy metal could predict the bacterial composition, we assessed the significance of the correlation using the “adonis2” function in vegan (55) (v 3.6.0). The relationship between heavy metals and *intl1* gene abundance we tested by a Spearman correlation.

### 2.9 Data availability

The sequence data (Microbial community and *intl1* amplicon data) is deposited at ENA (https://www.ebi.ac.uk/ena) under the accession number: PRJEB41986. The underlying ASV and metadata table can be found in the Supplementary Material (Water_Runoff_ASV_Table.csv,Water_Runoff_Metadata.csv,Sand_Filters_ASV_Table.csv, Sand_Filters_Metadata.csv).

## 3. RESULTS

### 3.1 Physico-chemical properties of road runoff, effluent of the SQIDs and filter media

As already described for this site by Helmreich et al. (56), the concentrations of Cr, Cu, Ni, Pb, and Zn as well as the pollution level (WPI_GFS_) of the analyzed road runoff (influent, I) and effluent samples of the two SQIDs (ESA, EF) showed strong seasonal variation with higher values observed in spring (Table 1). The higher EC in the spring samples indicate the influence of de-icing salt (sodium chloride) applied on-site, which contribute significantly to the toxicity of road runoff (33). As a consequence of the neutral to slightly alkaline pH of the samples, heavy metals were predominantly found in the particulate phase in the influent of the SQIDs. The dissolved Pb concentrations were below LOQ, as were half of the dissolved Cr and Ni concentrations. Consequently, it was only possible to determine the dissolved fractions of Cu and Zn, which were in median 18 and 21%. The overall pollution level, as indicated by the WPI_GFS_, of the SQID effluents was lowered with lowest total heavy metal contamination in EF. In ESA 18% of Cu and 38% of Zn were dissolved. In EF larger dissolved fractions were observed: 63% Cu and 40% Zn. In the filter material sample FD1 showed higher Ni contents than in FD2, but showed lower values for the residual metals, respectively.

### 3.2 Microbial parameters of road runoff and SQID systems

The investigated SQIDs were colonized by a diverse range of microbial taxa. About 7,538 unique amplicon sequence variants (ASV) were detected for water samples (I, ESA, EF) and 5,599 in filter material FD1 and FD2 (Table 2). The 16S copies as an approximation for cell counts ranged in the order of 10^8^-10^9^ copies per ml water. The copy numbers in the filter material was in the range of 10^7^ copies per gram material. The class I integron gene cassette *intl1* copy numbers had high numbers in both filter material and water samples. Neither strong seasonal effects nor differences between the filter materials in terms of 16S copies were detected.

**Table 2.**
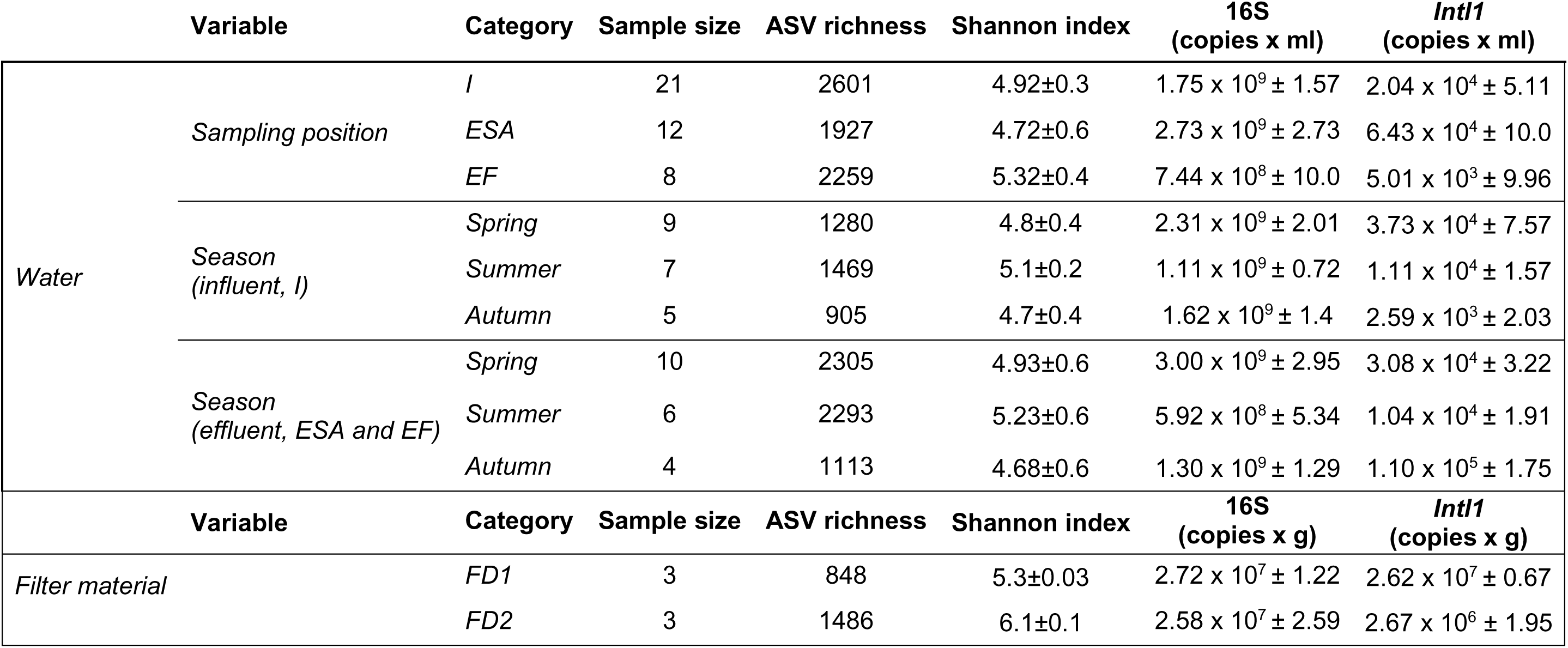
Water and filter media samples characteristics and statistical analysis for the different samples. I: Influent; ESA: Effluent of device D1; FD1: Filter material of D1; FD2: filter material of D2; EF: Effluent of device D2.

### 3.3 Microbial taxa of road runoff

In both systems, the most prevalent phyla consisted of Proteobacteria*, –* mainly composed by Gammaproteobacteria and Alphaproteobacteria, followed by Actinobacteriota, and Bacteroidota (Fig.2A). The main difference between the two devices was an increased proportion of Campilobacteriota for D1 that had itself established in the intermediate ESA (7%). At the genus level many genera ranged below 2% relative abundance (Fig. 2B). Most of the dominant genera like *Massilia*, *Alkanindiges*, *Sphingomonas*, *Hymenobacter, Acidovorax* and *Arthrobacter* that were found in the influent were still present in ESA. The ESA water samples showed a dominance of *Pseudarcobacter* (8%). In contrast to the water samples, the filter media of the SQIDs were clearly distinct (with minor vertical changes between the filter horizons; Figure S2). For both filter media, the most prevalent phyla consisted of Gammaproteobacteria and Alphaproteobacteria, followed by Actinobacteriota, Bacteroidota, Acidobacteriota, Chloroflexi, Desulfobacterota and Firmicutes (Fig.2A). On the genus level, *Hydrogenophaga* and *Rhodoferax* (4.7% and 4.1%, respectively) were the dominant *taxa* in FD1 column, while *Arenimonas* (3.4%) and *Sphingomonas* (2.8%) dominated FD2 (Fig. 2B)

**Figure 1.**
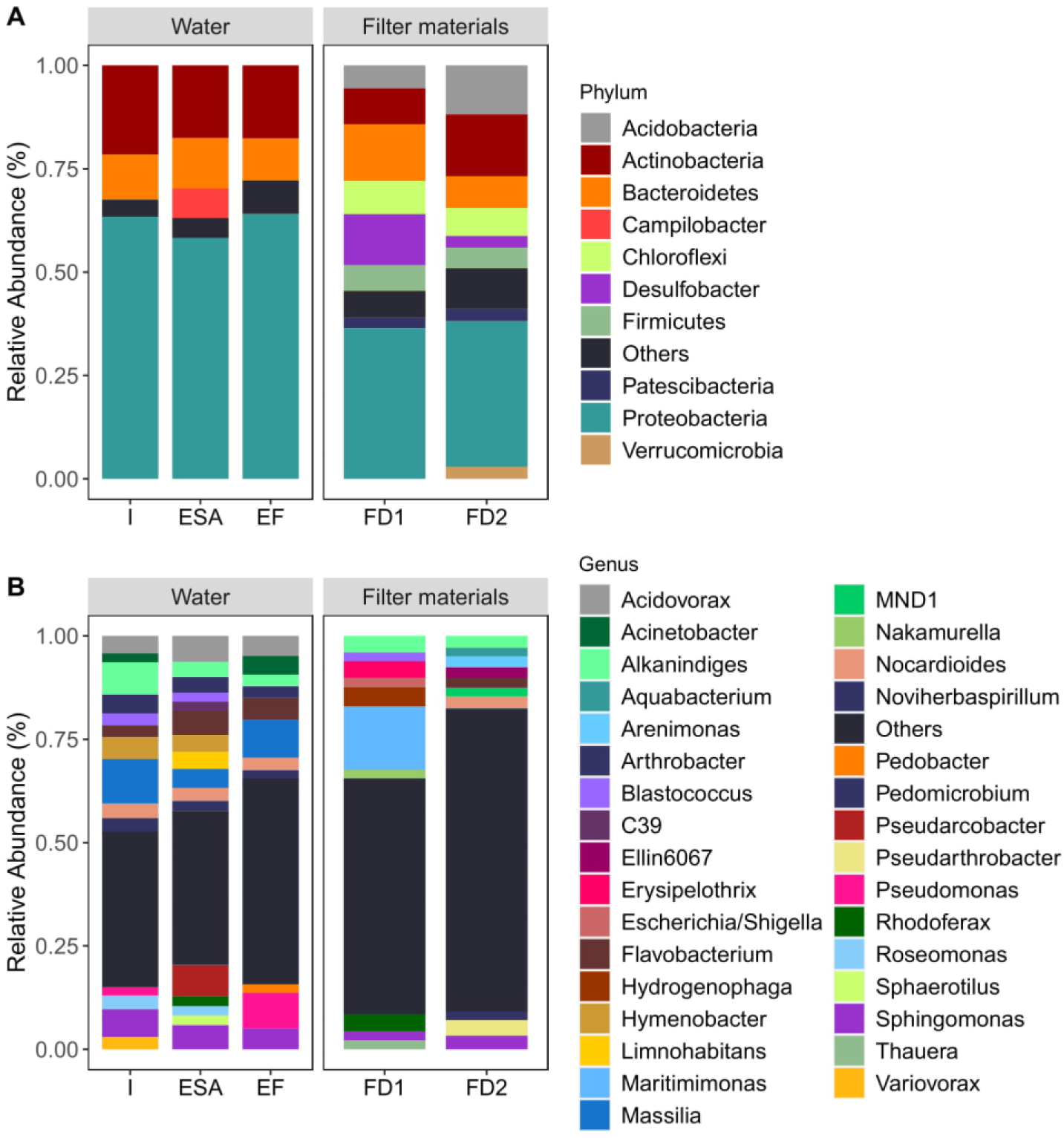
Distribution bar plot of the relative abundance of bacterial groups at Phylum (A) and Genus (B) level in untreated and treated road runoff and SQIDs’ filter media. For better representation only taxa with relative abundance > 2% are displayed. I: Influent, ES: effluent of sedimentation, SA: effluent of sedimentation and adsorption, EF: effluent of filtration; FD1: filter material of D1; FD2: filter material of D2 (D1 and D2 as depicted in Figure 1.)

**Figure 2.**
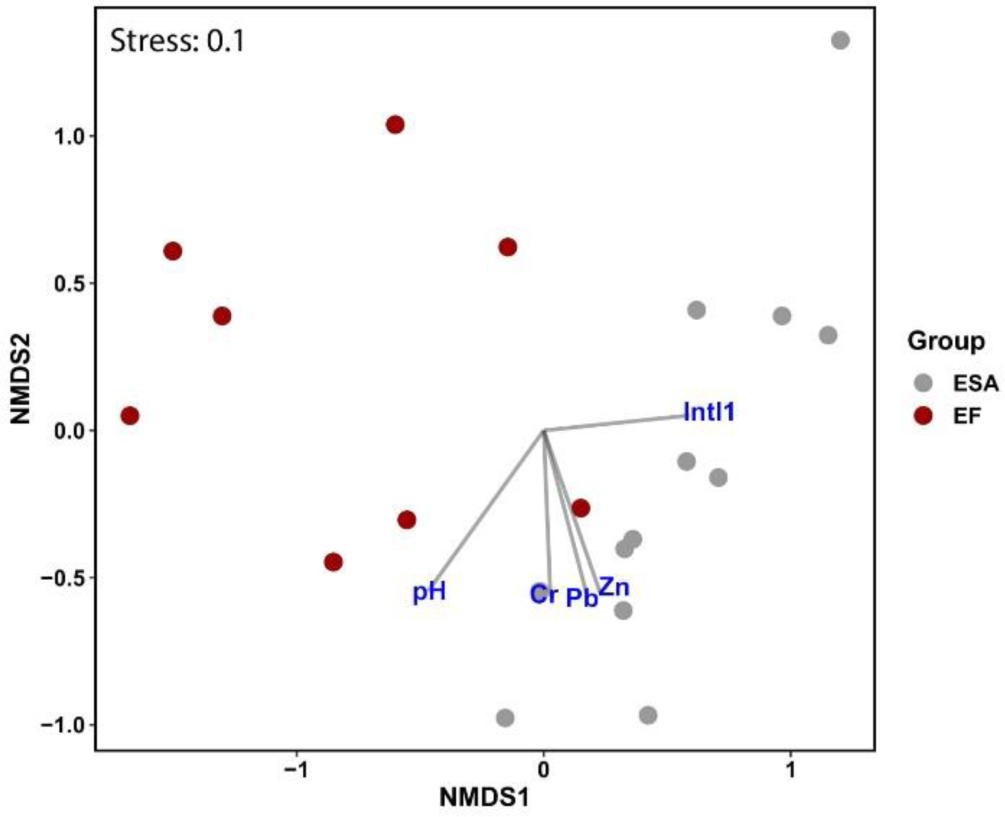
Non-metric multidimensional scaling (NMDS) based on Bray-Curtis dissimilarity of SQIDs effluents community data (n=20) and environmental factors. ESA: effluent of sedimentation and adsorption, EF: effluent of filtration. The correlation between species and environmental variables are indicated by a perpendicular projection of the species arrow-tips onto the line overlaying the environmental variable arrow.

### 3.4. Retention of microbes by SQIDs

By partitioning the β-diversity into loss of species (nestedness) and species turnover, we could confirm that we mainly see a turnover of taxa (as ASV) between influent and effluent of D1 (turnover = 0.81, nestedness = 0.04) and D2 (turnover = 0.90, nestedness = 0.02), pointing to rather a replacement of species along the water’s flow of the SQIDs, than a species loss along the environmental gradient (Overall nestedness = 0.03). Differential abundances of microbial genera pointed to few differentially enriched genera for D1 and D2 (Figure S3). In D1 few genera showed up, but a high enrichment of C39 (Rhodocyclaceae; log2 fold change of 28.5) was observed. In D2, a stronger removal was detected with 15 different genera with up to 21.2 log2 fold change. On the ASV level, we identified several potential microbial risk factors, i.e., taxa that are derived from animal host systems and may be relevant for human health and hygiene (e.g. *Erysipelothrix*, *Shigella*, *Escherichia*, Table S2). The majority of these taxa were mostly found at very low relative abundances in the road runoff (<0.08%). Among the potential bacterial pathogens, the genus *Pseudomonas* was dominant in all samples followed by *Corynebacterium* in the effluent ESA.

### 3.5 Factors that influence the microbial community composition

Multivariate statistics separates the two effluent water samples EF and ESA, and further point to single metals, pH, and *intl1* as additional influencing factors (Figure 2, Table S3). Moreover, seasonal changes and heavy metals (as sum of Cr, Cu, Ni, Pb and Zn molarity), impacted the species composition of the effluent samples (GLM: LRT = 12395, LRT = 9350, p = 0.006 and p = 0.03, respectively).

**Table 3.** Kruskall-Wallis test employed to identify statistical differences in bacterial richness (ASV), diversity (Shannon), 16S and Intl1 gene copies along the two SQIDs water treatment. P-value significance codes: < 0.001 ***; < 0.01 **; < 0.05 *. n.a. (not applicable).

|  | n | Shannon |  | 16S<br>log(16S gene copies/mL) |  | *Int1*<br>log(int1 copies/mL) |  |
| --- | --- | --- | --- | --- | --- | --- | --- |
|  |  | Chi-squared | increased in | Chi-squared | increased in | Chi-squared | Increased in |
| D1 Influent vs effluent | 24 | 1.76 | n.a. | 0.65 | n.a. | 3.88* | Effluent (+2.01) |
| D2 Influent vs effluent | 17 | 4.28* | Effluent (+1.09) | 3 | n.a. | 0.85 | n.a. |

### 3.6 The influence of heavy metals on microbial taxa

In order to gain more insight on the role of heavy metals, we preselected the most predictive metals using bioenv, which indicated an influence of Ni, Zn and Cu on the microbial community (adonis2 for the total model: R^2^ = 0.28, p < 0.001). Likewise, the 16S normalized *intl1* abundance showed linear relationships with the heavy metals concentrations Ni, Pb, and Zn with higher explanatory power for Ni and Zn (R^2^ > 0.71, p < 0.001; Figure 3). This was further explored by co-correlating the most abundant 50 genera with, heavy metals, and *intl1* (Figure 4). A total of seventeen genera showed positive correlations with the metal concentrations, with *Aquabacterium, Hydrogenophaga* and *Trichococcus* associated with almost all the measured metals. Three out of the five heavy metals (Ni, Pb and Zn) showed the highest positive association with the relative abundance of genera (R^2^ > 50, p < 0.05). On the other hand, *Aeromonas*, *Aquicella, Legionella* and *Pseudomonas* were showing significantly negative correlations to heavy metals.

**Figure 3.**
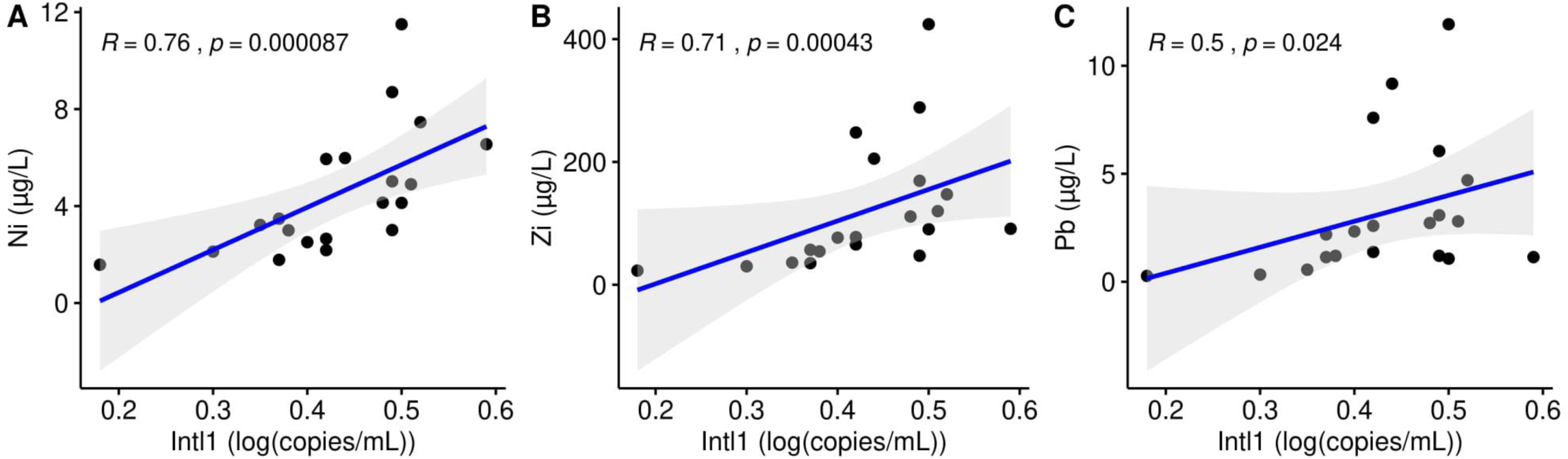
Spearman correlations between biomass normalized *intl1* and heavy metals in SQIDs effluent water samples (n=20, nickel (A), zinc (B), lead (C)).

**Figure 4.**
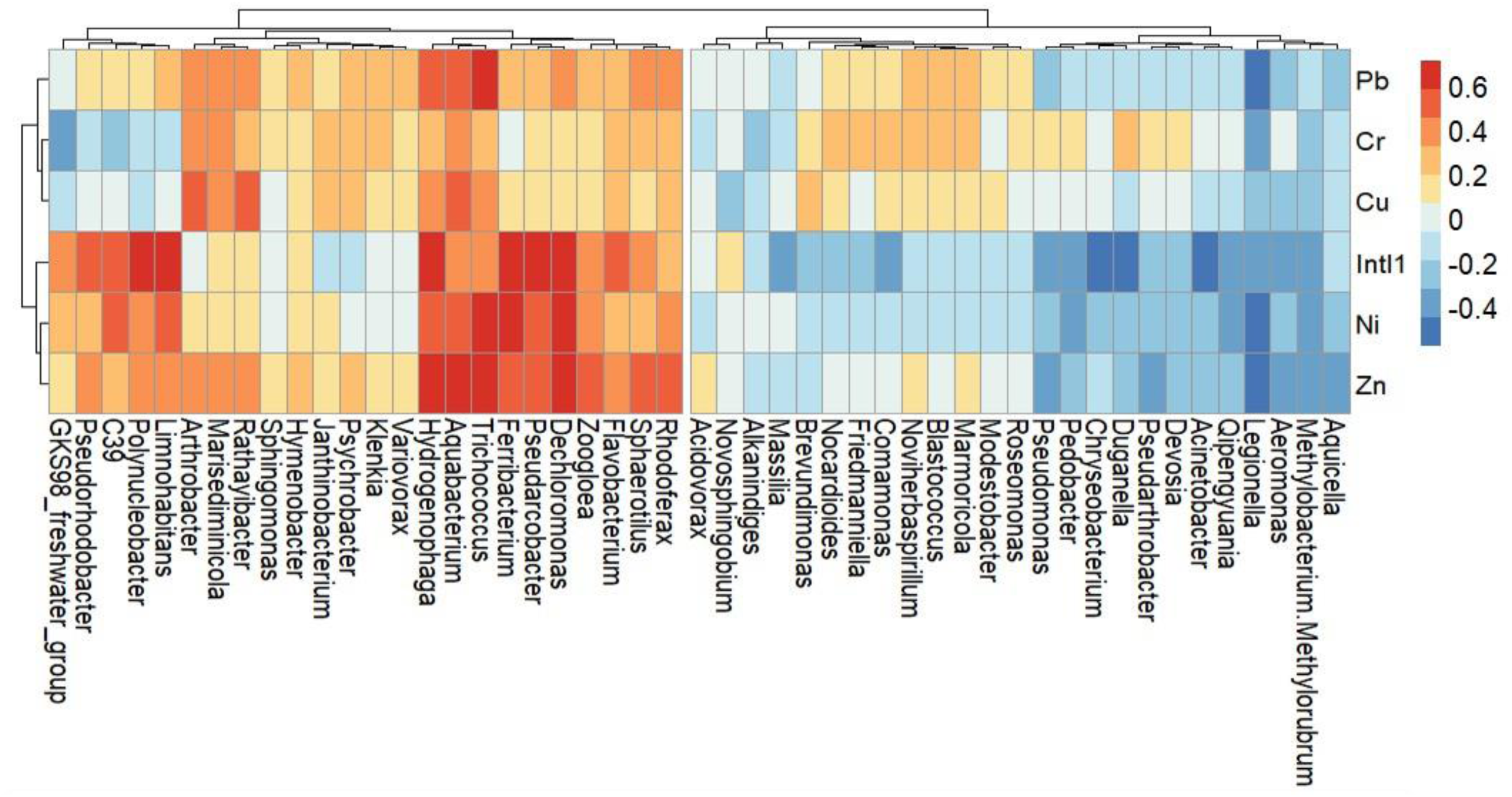
Heat map of Spearman correlation analysis between relative abundance of water effluents (n = 20) bacterial community and content of heavy metals at Genus level. Colors depict individual negative and positive correlations.

### 3.7 The role of *intl1* in facilitating heavy metal resistances

Eleven genera were also co-correlating with *intl1* (*C39*, *Dechloromonas*, *Ferribacterium*, *Flavobacterium, Hydrogenophaga, Limnohabitans, Polynucleobacter, Pseudarcobacter, Trichococcus and Zoogloea,* R^2^ > 0.45 p < 0.05). To further test these potential linkages of *intl1* and heavy metal resistance, we sequenced parts of the genes that were carried by the class 1 integrons in the systems. In total, 296 of the 622,526 reads from the integrons (0.05%) contained 98 different resistance genes. Of these, 82 were metal or biocide resistance genes (BacMet), 7 were clinically relevant antibiotic resistance genes (ResFinder) and 11 were antibiotic resistance genes previously only encountered in functional metagenomics studies (FARME). The most common antibiotic resistance genes were aminoglycoside resistance genes *aadA5* (found on 6 integron sequences) and *aadA4* (3 sequences), and fluoroquinolone efflux pump *oqxB* (4 sequences). Four other genes (*aac*(*3*)*-Ia*, *aac(3)-Ib*, *msr(D)* and *vat(E)*) were found only once. Most of the identified genes were involved in metal resistance, most commonly to heavy metals such as Pb, Cd and Zn (Figure 5). Cu and Ag resistance genes constituted around 13% of the identified genes, while antibiotic resistance genes accounted for 8.6% of the identified genes in total. Biocide resistance genes made up approximately one-third of the identified resistance genes. The most commonly encountered resistance genes (> 6 occurrences) were the metal resistance genes czrA, czcA and silP, the biocide resistance gene qacE, and the efflux pumps mexW and mexI that also facilitate multidrug resistance (Table S4).

**Figure 5.**
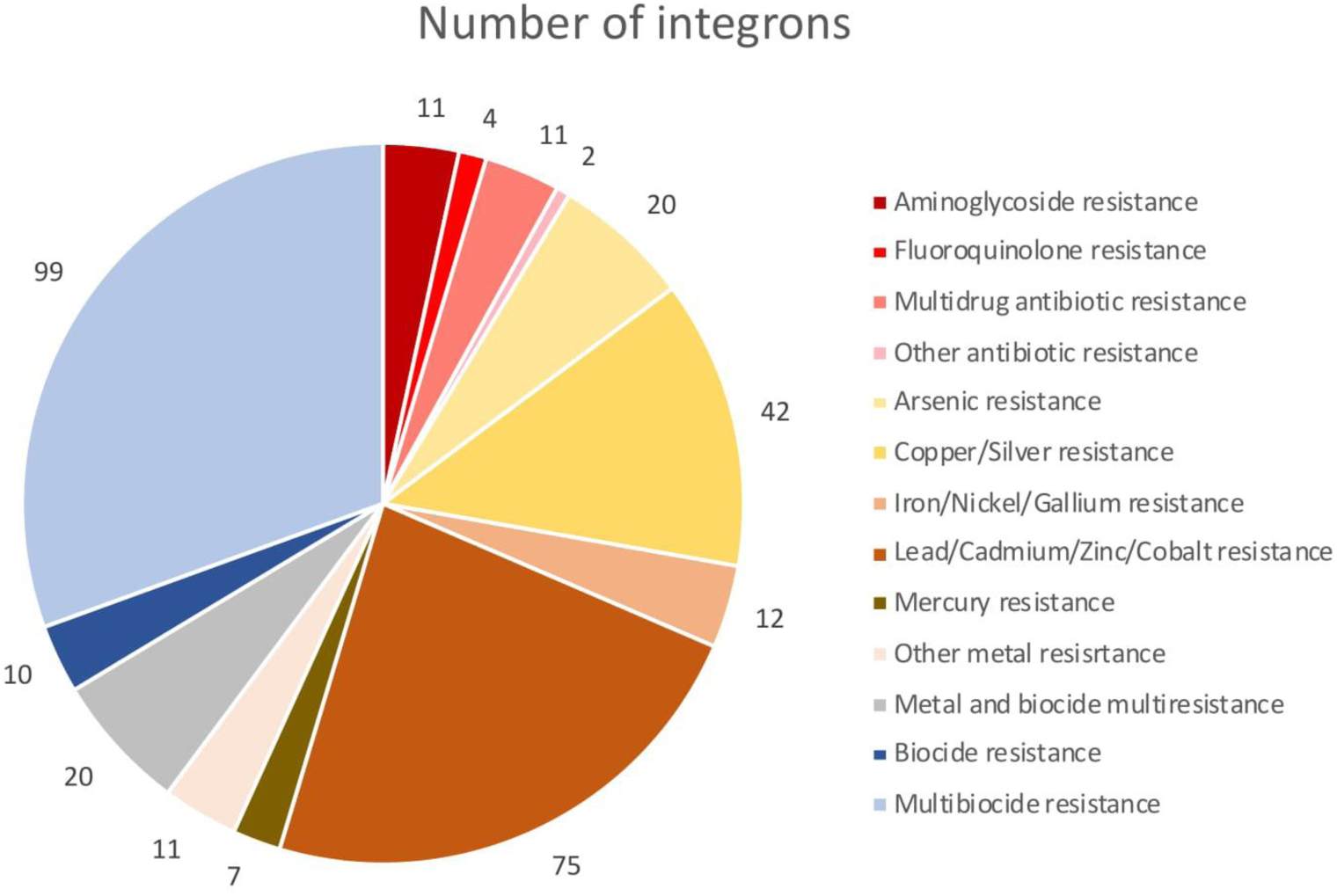
Distribution of different types of resistance genes in the class I integron gene cassette sequences

## 4. DISCUSSION

The results revealed that SQIDs not only retain heavy metals from road runoff, but also change the microbial community composition, alter the microbial load, and influence the mobile genetic elements. The overall analysis of the road runoff and the SQID samples indicated a predominance of Gammaproteobacteria, Actinobacteriota and Bacteroidota in the water and the respective filters media. These findings are consistent with previous reports, where these phyla have been identified in stormwater runoff as anthropogenic or erosion signatures (57–59). Similarly, the taxa that occurred in the SQID filter materials, like Desulfobacterota, Chloroflexi and Acidobacteriota, were all previously observed in an infiltration basin collecting highway runoff (17). Acidobacteriota are mainly found in low pH environments (60) tolerating various pollutants such as PCBs, petroleum compounds (61, 62) and heavy metals (63). Only few taxa with pathogenic potential were present at low levels, and SQIDs are not designed for microbial retention, but their occurrence in the road runoffs warrants further investigation, in particular when the water treatment selects for mobile genetic elements and when the receiving waters are considered as critical resource.

### 4.1 Influence of heavy metals on the microbial community

Heavy metals with high concentrations in waters or soils show toxic effects to almost all microbes by affecting metabolic functions such as protein synthesis (64, 65), thus leading to variations in microbial biomass and diversity (66). Several studies have shown how Cu, Zn, Pb and other heavy metals severely inhibited microbial biomass and could cause a reduction of microbial α-diversity (67). The most common conclusion is that only high concentrations can significantly decrease bacterial biomass, whereas mid-low concentrations of heavy metals can increase microbial biomass and stimulate microbial growth (68, 69). Our pH and metal measurements indicated that large fractions of the heavy metals are not readily bioavailable, nevertheless particulate-bound heavy metals are considered partly bioavailable (70). Anoxic conditions in particular may favor metal reduction as a source of energy, which generally leads to the release of metal ions into the water (e.g.,Teiri et al., 2016). The prevalence of Desulfobacterota, which are responsible for sulfate reduction processes in stormwater retentions ponds (72), together with Chloroflexi that constitute a substantial proportion of the activated sludge community in wastewater treatment plants (73), point to anoxic processes that may occur in the SQID systems.

### 4.2 Heavy metal resistances are linked to *intl1*

Ni and Zn, that showed and influence on the microbial community composition are known to induce different resistance mechanisms in bacterial metabolism (74–76). In this context, one interesting case was Arcobacter (and *Pseudoarcobacter*; Pérez-Cataluña et al., 2018), which was abundant in water and filter material. *Arcobacter* is known to form biofilms in various pipe surfaces, such as stainless steel, Cu, and plastic, colonizing water distribution systems (78, 79). In our case, *Pseudoarcobacter* mainly correlating to Ni and Zn also,showed a strong correlation with *intl1* gene abundance, thus suggesting the selection for bacterial resistance in *Pseudoarcobacter*. Both, metal and antibiotic resistance are commonly carried on mobile genetic elements. Integrons, in particular, have been recognized as marker for anthropogenic pollution (80). Prior research from different heavy metal polluted scenarios showed the development of resistances due to horizontal gene transfer (81, 82), and there have been signs of co-selection of several resistant genes linked to clinically relevant antibiotic resistance (83). For example, resistance to As, Mn, Co, Cu, Ag, Zn, Ciprofloxacin, β-lactams, chloramphenicol and tetracycline is achieved by reduction in membrane permeability (84, 85). Similarly Cu, Co, Zn, Cd, tetracycline, chloramphenicol and β-lactams resistance is achieved through rapid efflux of metal or antibiotic (86, 87). Therefore, heavy metals have potential to represent extended selection pressure for development of antibiotic resistance in microorganisms (88), and the transfer of these resistant bacteria in the environment may pose potential risks to human health (89).

### 4.3 Stormwater quality improvement devices as hot spots for horizontal gene transfer

The *intl1* gene cassette analysis highlighted the presence of heavy metal resistances in these microbial communities, and the abundance of *intl1* in the filter media, compared to the influent, suggests a strong selection pressure that aligns with a significant rate of horizontal gene transfer taking place in the systems. The amount of bacteria carrying class 1 integrons is consistent with data reported in polluted water systems like WWTPs (90). However, while previous research, normally described high removal rates of bacteria carrying class 1 integrons (55%) after treatment process (90), consistent with our findings, few studies has reported an increase in the abundance of the *intI1* gene during the wastewater treatment (91, 92). The variation of results observed among studies may be attributed to several factors such as selected resistant bacterial taxa, the climat and population conditions, occurrence of rain events before sampling as well as organic loading, pH and temperature. Horizontal gene transfer plays an important role in the evolution, diversity and recombination of multi-drug resistant strains (93, 94). The class 1 integron has been associated with the presence of metal resistance genes (MRGs) and antibiotic resistance genes (ARGs) (95, 96). Our data suggests that SQIDs could be a high-risk environment for resistance development, similar to other hotspots, like manure, sewage and municipal solid waste (97–99). Furthermore, the presence of different resistance genes (e.g. *czrA*, *czcA* and *silP*), including the high proportion of multidrug resistance genes (e.g. *mexW*, (100, 101)) as well as the strong positive correlations to heavy metals, suggest that integrons contribute to the spread of MRGs and ARGs within road runoff drainage systems. While no typical antibiotic treatment related resistance genes (*sul1*, *ampC*, etc.) were identified, they may be present on other mobile genetic elements not targeted in this study.

### 4.4 Limitations and future directions

This study provides a first deeper description of road runoff microbial communities and contributes to our understanding of their potential environmental impact on the receiving water bodies. As a pioneer study, our study design was limited to two SQIDs and we could only monitor three seasons. Thus, we could not investigate if there are further effects by e.g. higher amounts of de-icing salts in winter, which potentially enhance mobility and bioavailability of the present heavy metals (33,102,103). Furthermore, we did not consider other systems such as infiltration basins or sand filters, and it remains an open question if our results are transferable to other road runoff drainage systems. However, it is obvious that runoff from a highly trafficked urban road carries a high microbial load with dominant signs of anthropogenic pollution. This comes with a relatively high risk related to the cycling of resistance genes and thus microbial risk mitigation practices should be considered in the future. Recently, it has become clear that microbes are critically linked to our changing environment, and that they have to be included in future policies (104). Future studies are therefore encouraged to assess the risks of discharge of microbes and their resistance genes from SQIDs and other SCMs into receiving environments.

## Supporting information

Water_Runoff_ASV_Table

Water_Runoff_Metadata

Sand_Filters_ASV_Table

Sand_Filters_Metadata

## 5. ACKNOWLEDGEMENT

We would like to thank Heidrun Mayrhofer and Hubert Moosrainer for technical assistance. RL acknowledges the International PhD Programme “Environment, Resources and Sustainable Development” scholarship (#Parthenope University of Naples), monitoring of the SQIDs by BH and SR was funded by the Bavarian Environment Agency (AZ: 67-0270-96505/2016 and AZ: 67-0270-25598/2019). JBP was supported by the Sahlgrenska Academy at the University of Gothenburg, the Swedish Research Council (VR; grant 2019-00299) under the frame of JPI AMR (EMBARK; JPIAMR2019-109), and the Centre for Antibiotic Resistance Research at the University of Gothenburg (CARe). CW was financially supported by TUM Junior Fellowship and the DFG project (WU 890/2-1). We would like to thank Dr. Klaus Neuhaus of the ZIEL Core Facility Microbiome, Technical University of Munich, Freising, Germany, for sequencing.

## Supplementary data

**Table S1.** Insignificance threshold values for groundwater specified by the Länderarbeitsgemeinschaft Wasser (LAWA) in µg/L, which were used to calculate the water pollution index (WPI_GFS_)

| Cr | Cu | Ni | Pb | Zn |
| --- | --- | --- | --- | --- |
| 3.4 | 5.4 | 7.0 | 1.2 | 60.0 |

**Table S2.** List of potential pathogenic taxa identified in road runoff and effluent of stormwater quality improvement devices. ESA: effluent of sedimentation and adsorption, EF: effluent of filtration. All the values are reported as percentages.

| Genus | Influent | ESA | EF |
| --- | --- | --- | --- |
| Aeromonas | 0.07 | 0.08 | 0.58 |
| Burkholderia-Caballeronia-Paraburkholderia | 0.01 | 0.03 | 0.10 |
| Clostridium_sensu_stricto_1 | 0.08 | 0.03 | 0.01 |
| Corynebacterium | 0.03 | 0.16 | 0.05 |
| Cryptosporangium | - | 0.005 | - |
| Enterobacter | 0.004 | 0.001 | - |
| Erysipelothrix | 0.015 | 0.04 | - |
| Enterococcus | - | - | 0.006 |
| Escherichia/Shigella | 0.001 | - | - |
| Legionella | 0.03 | 0.08 | 1.91 |
| Mycobacterium | 0.04 | 0.06 | 0.04 |
| Pseudomonas | 2.04 | 0.47 | 8.64 |
| Serratia | 0.01 | 0.007 | 0.009 |
| Staphylococcus | - | - | 0.05 |
| Yersinia | 0.001 | 0.003 | 0.02 |

**Table S3.**
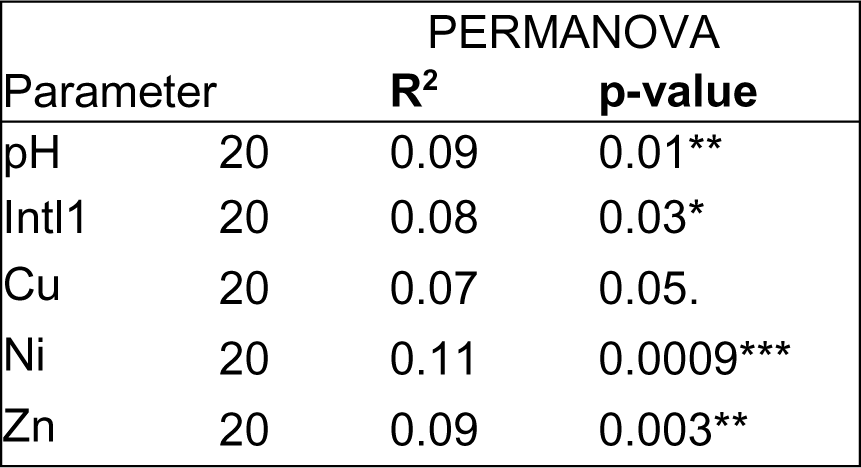
Results from PERMANOVA analysis to test the effects of environmental parameters on microbial community composition treated runoff by the two SQIDs (effluents, n=20). p-value significance codes: < 0.001 ‘***’; < 0.01 ‘**’; < 0.05 ‘*’, 0.05 “.”.

**Table S4.** Resistance genes encountered on at least six of the sequenced integrons

| Gene | # integron sequences | Function | Database |
| --- | --- | --- | --- |
| *czrA* | 24 | Zn and Cd resistance regulator | BacMet |
| *qacE* | 20 | Quaternary ammonium compound-resistance (exporter) | BacMet |
| *czcA* | 20 | Cd, Zn and Co resistance (exporter) | BacMet |
| *mexW* | 14 | Biocide exporter | BacMet |
| *silP* | 13 | Ag efflux system periplasmic chaperone | BacMet |
| *arsB* | 10 | As Sb exporter | BacMet |
| *vexB* | 9 | Multidrug efflux transporter (biocide resistance) | BacMet |
| *merA* | 7 | Hg reductase enzyme | BacMet |
| *acrB* | 7 | Multidrug efflux protein (substrates include biocides as well as the antibiotics ampicillin and chloramphenicol) | BacMet |
| *vexH* | 6 | Multidrug efflux protein (including antibiotics ampicillin and novobiocin) | BacMet |
| *recG* | 6 | DNA repair protein, involved in Cr, Te and Se resistance | BacMet |
| *nia* | 6 | Cation transport ARPase, involved in Fe and Ni resistance | BacMet |
| *adeE* | 6 | Multidrug efflux transporter (biocide resistance) | BacMet |
| *aadA5* | 6 | aminoglycoside nucleotidyltransferase | ResFinder |

**Figure S1.**
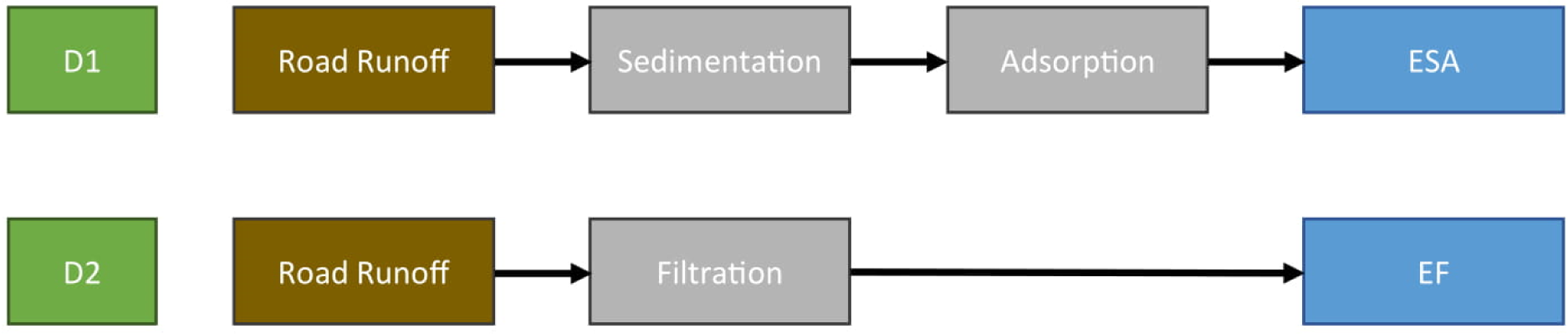
Structure and treatment processes of the monitored SQIDs. Name of the two SQIDs are shown in green (D1 and D2), main processes are showed in gray boxes, effluents classification are in blue boxes. ESA: Effluent of Sedimentation and Adsorption, EF: Effluent of Filtration. The arrows depict the flow of the water.

**Figure S2.**
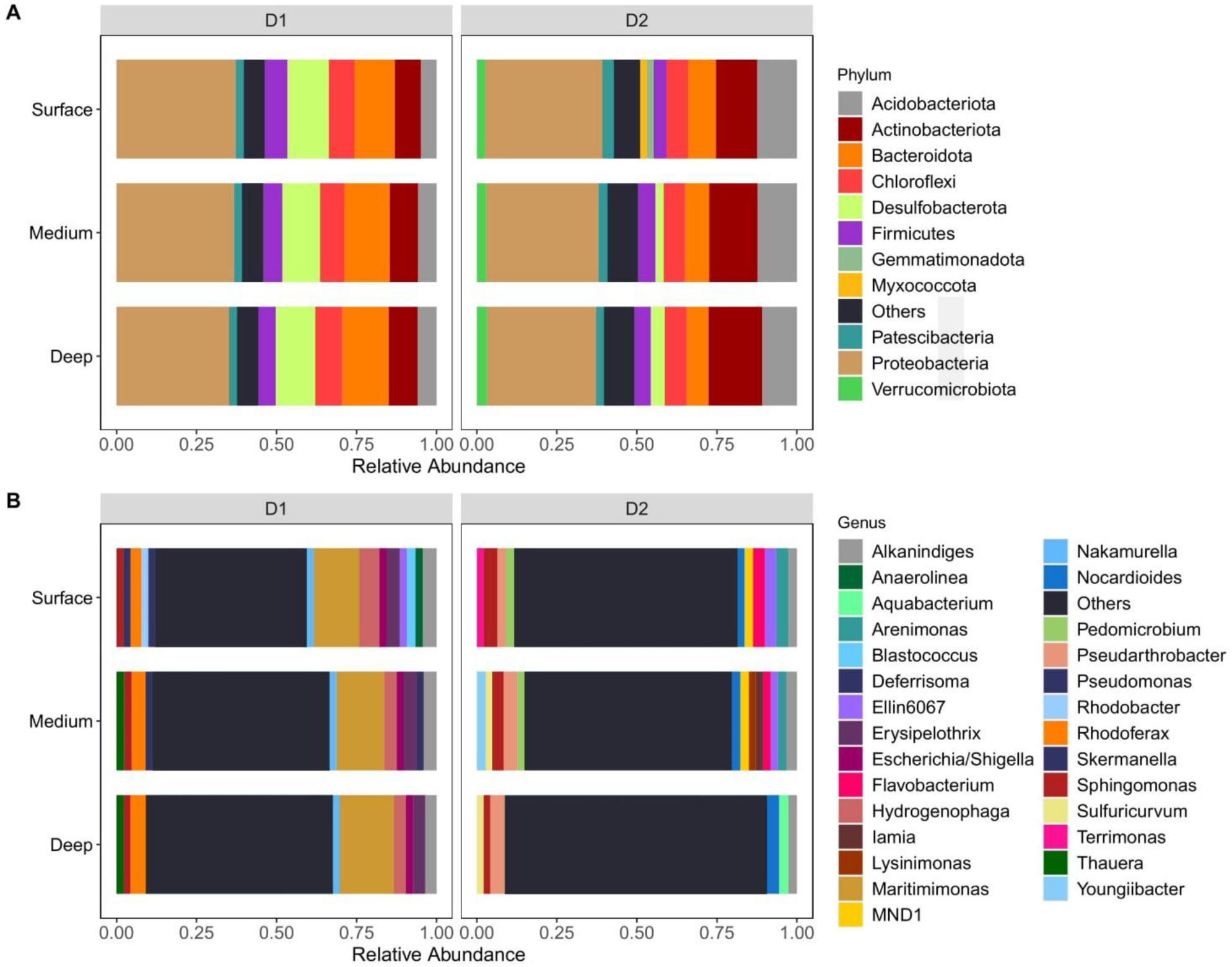
Distribution bar plot of the relative abundance of bacteria community at Phylum (A) and Genus (B) level along different depth layers of filter media samples. Surface (0-5 cm), Medium (5-10 cm), Deep (10-15 cm). For the best representation only taxa with relative abundance > 2% are displayed.

**Figure S3.**
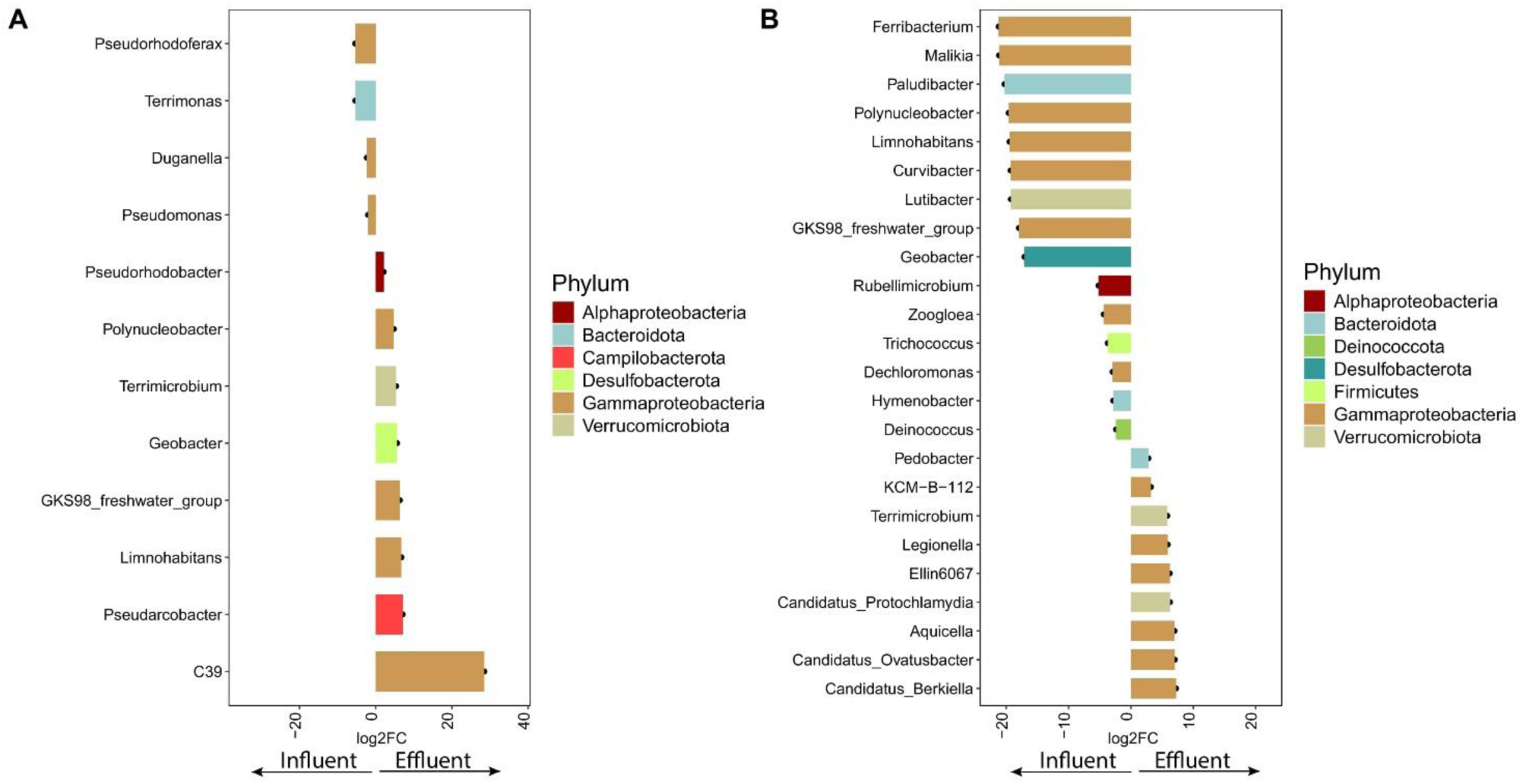
Differential abundance analysis of bacterial relative abundance in D1 (A) and in D2 (B) effluents (n=20) compared to those of the influent. Core taxa with statistically significant difference (log2FC > ± 2; adjusted p-value < 0.05) are displayed in the plot. The different colours depict bacterial phyla. Red arrow indicate taxa present in the influent, black arrow show taxa in the effluents.

